# Psilocybin lengthens hippocampal sharp wave ripples

**DOI:** 10.64898/2026.08.17.745042

**Authors:** Ilknur Kayikcioglu Bozkir, Ryan Lashin, Tiecheng Liu, Dinesh Pal, Kamran Diba, Nathaniel Reid Kinsky

**Affiliations:** Gumushane University; University of Michigan; University of Michigan Medical School

## Abstract

Psilocybin is a psychedelic which has been shown to induce neural plasticity through activation of intracellular serotonergic 5-HT2A receptors. It also produces brain-wide changes in structural and functional connectivity and holds promise as a therapeutic compound for treating anxiety and depression. Despite links between psilocybin-induced plasticity, the psychedelic experience, and reduction in depressive symptoms, little is known about the effects of psilocybin on the function of the highly plastic hippocampus, a region crucial for memory whose dysfunction is linked to neural disorders such as depression and anxiety. In this study, we investigated the acute and lasting effects of psilocybin on rodent sharp-wave ripples (SWRs), transient high frequency oscillations observable in the hippocampal local field potential which are linked to memory consolidation. We found that a 10 mg/kg dose of psilocybin robustly decreased the peak SWR frequency and increased the duration of SWRs immediately following administration compared to control sessions the day before and after. Psilocybin also perturbed sleep architecture, resulting in a pronounced reduction in non-rapid eye movement (NREM) sleep which lasted for hours. Therefore, psilocybin could impact memory processing by modulating hippocampal SWRs.

## Introduction

Psilocybin is a naturally occurring psychedelic compound with growing evidence of therapeutic effects in the treatment of depression and anxiety^1-4^. Psilocybin’s psychedelic and therapeutic effects are mediated primarily through agonism at serotonergic 5-HT2A receptors^5^. A previous study demonstrated that psilocybin can promote both structural and functional neuroplasticity in the frontal cortex^6^, a region strongly linked to major depressive disorder^7^, and that this plasticity enabled the frontal cortex to re-organize brain-wide structural and functional connectivity^8^.

Besides the frontal cortex, the hippocampus is one potential hub which could mediate psilocybin’s therapeutic effects. Reduced hippocampal volume has been linked with unipolar depression^9-11^, and patients with major depressive disorder frequently exhibit cognitive deficits in hippocampal-dependent memory tasks^12,13^. Moreover, reversal of volumetric degradation in the dentate gyrus subregion of the hippocampus by depression has been linked to improvements in major depressive disorder symptoms^9,14,15^. The hippocampus is highly plastic^16,17^ and strongly innervates a wide range of cortical and subcortical regions^18,19^. Like the prefrontal cortex, it is rich with 5-HT2A receptors^20^ which promote plasticity^21-23^ and are important for learning and memory^24^. Additionally, hippocampal activation can reduce stress-related physiological responses due through inhibition of the hypothalamic-pituitary axis^25^. Thus, the hippocampus is a strong candidate for inducing psilocybin-related plasticity across the brain.

Hippocampal sharp wave ripples (SWRs) are high-frequency oscillations observed in the hippocampal region CA1 local field potential during periods of awake rest and sleep. SWRs consist of a large amplitude 1-4Hz deflection detected in the CA1 *stratum radiatum* called the *sharp wave* and a transient 150-250Hz oscillation localized to the CA1 pyramidal cell layer called the *ripple*^26^. SWRs coincide with bursts of spiking activity called replays which closely match the patterns of spiking observed during active behavior in a time compressed manner^27-29^. Sleep SWRs are causally linked to memory consolidation^30,31^, presumably by coordinating information flow between CA1 and cortical regions^26^. Awake SWRs are tied to learning^32^ and could support planning during memory-guided tasks through awake replay of past and future trajectories^33,34^. However, recent studies indicate that awake SWRs and associated replays may instead subserve long term memory storage by rehearsing recently learned information^35,36^ and prioritizing memories for later replay during sleep^37^.

SWR features provide information regarding hippocampal circuit function. First, sharp waves co-occur with population burst spiking events in upstream CA3^38^; thus, the sharp wave amplitude reflects the strength of the connection between CA3 and CA1^39^. Second, ripple frequency is modulated by interactions between excitatory pyramidal cells and interneurons localized near *stratum pyramidale^40^*. Finally, ripple duration is linked to memory load and novelty, and artificially lengthening SWR duration enhances memory strength^41^. While psilocybin has been reported to have widespread effects on cortical oscillations and functional connectivity in rodents^1,42^, little is known about its effects on hippocampal function. Here, we investigate acute and lasting effects of psilocybin administration on hippocampal sharp wave ripple features to understand how psilocybin impacts the hippocampal circuit.

## Results

Following chronic implant of a linear silicon probe in the intermediate hippocampus and v4 UCLA miniscope over the prefrontal cortex (data not shown here), 5-12 month old Long Evans rats (n= 2 male and 2 female) underwent three recording sessions over 3-4 days all of which occurred in a rest box similar to the animal’s home cage (**Figure 1A**). In the first session (SALINE1), neural activity was recorded for ∼1 hr following an intraperitoneal (i.p.) injection of saline to establish a baseline response to the experimental procedure. The second session (PSILOCYBIN) occurred 1-2 days later and followed the same general procedure except that animals were given a short baseline recording period (PRE, ∼5-15 min) following which they received a 10 mg/kg i.p. psilocybin injection and the subsequent recording (POST) was lengthened to 3-4 hours to capture the acute effects of psilocybin. The following day, animals underwent the same procedure as the first day (SALINE2) to capture any persistent effects of psilocybin administration. Two rats also had a PRE recording for the Saline1/Saline2 sessions, and SALINE1/SALINE2 recording durations were also extended to 3-4 hours to match the duration of the PSILOCYBIN recording in one rat. See **Figure 1A** for the full experimental schedule. The dose for psilocybin was based on our recent studies^42-44^. Animal motion was captured using the UCLA Miniscope on-board head orientation sensor and through high-resolution videos and markerless tracking with DeepLabCut^45^.

**Figure 1:**
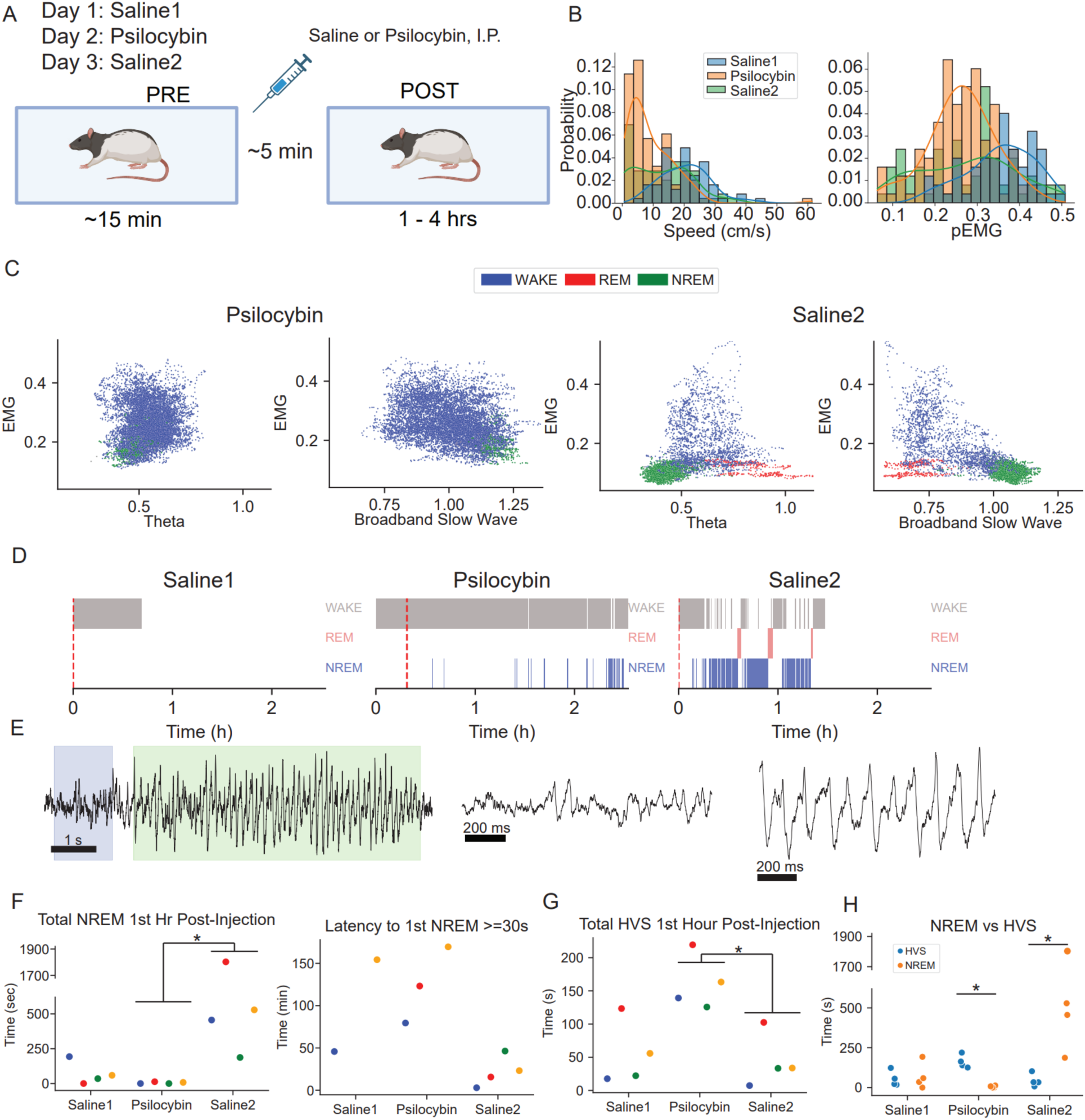
Experimental setup and effects on sleep architecture. **A)** Experimental schedule across three consecutive days. A typical recording session consisted of a short recording (PRE) prior to i.p. injection of saline or psilocybin, followed by a 1-4 hr POST injection recording. **B)** Histogram of speed (left) and pEMG (right) values across all sessions with kernel density estimates overlaid. **C)** Scatterplot of pEMG vs broadband slow wave power (left) and pEMG vs Theta power ratio (right) with dashed lines showing thresholds used to separate NREM from other states (vertical) and REM from wake (horizontal) from one rat. **D)** Example hypnogram from same rat as C demonstrating little to no sleep following psilocybin injection. Red dashed line: time of saline or psilocybin i.p. injection. **E)** (left) Example local field potential from electrode in *stratum pyramidale* showing example wake period (blue, middle) and HVS epoch (green, right). **F)** (left) Total time in NREM in the first hour after injection. (right) Latency to first extended NREM bout (right); note that Animal 3 did not enter NREM during the PSILOCYBIN session. Colors indicate different animals. **G)** Total HVS time in the first hour after injection. **H)** Total NREM vs HVS time in the first hour after injection. *p=0.029 two-sided Mann-Whitney U-test.

We first assessed locomotion following saline vs psilocybin injections. We observed a unimodal distribution of speed during SALINE1 (median=20.5 cm/s) and PSILOCYBIN (median=6.5 cm/s), and a bimodal distribution for SALINE2 with median=11.9 cm/s (**Figure 1B**, left). We next assessed changes to brain state by measuring three metrics: spectral power in the theta range divided by the overall spectral power (theta / all ratio), power spectral slope (PSS, a proxy for power in the broadband slow wave frequency range of 1-4 Hz), and pseudo electromyogram (pEMG) in 1 second bins^46^. We observed a bimodal distribution of pEMG in SALINE2; in contrast, we observed unimodal pEMG distributions during both SALINE1 (median=0.36) and PSILOCYBIN (median=0.27) with very few low pEMG values suggesting that little sleep occurred following injection in these sessions. To confirm this, we next identified periods of non-rapid eye movements sleep (NREM, characterized by high PSS) rapid eye movement sleep (REM, characterized by low PSS, high theta power, and low pEMG), and wake (remaining periods) using established methods^46^ (**Figure 1C-D, Supplemental Figure 1-Supplemental Figure 2**). Indeed, animals exhibited significantly less NREM during PSILOCYBIN compared to SALINE2 (**Figure 1F, left**) despite remaining highly immobile (**Figure 1B**, left). All animals also exhibited a longer latency to the first extended NREM bout during PSILOCYBIN compared to SALINE2 (**Figure 1F, right**). We observed high voltage spindles (HVS) – large amplitude low frequency deflections (**Figure 1E)** across all channels which typically emerge during wake-to-sleep transitions^47,48^ – during all sessions. However, we found the total time of HVS increased for all animals during PSILOCYBIN compared to SALINE1 and SALINE2 session (**Figure 1G**). Finally, we compared time spent in NREM vs HVS. During Saline1, we observed similar, low levels of both NREM and HVS, likely due to high levels of arousal following the first injection. However, time spent in HVS was significantly higher during PSILOCYBIN, while NREM time was significantly higher during SALINE2 (**Figure 1H**). Therefore, psilocybin disrupted locomotion and reduced normal sleep amount following drug administration^49^.

We next examined how psilocybin administration impacted hippocampal SWRs. SWRs were detected as epochs in the ripple-band (150-250 Hz) filtered signal whose power exceeded 4 standard deviations above the mean for at least 50ms. First, we observed no change in ripple rate across sessions (**Figure 2A).** We next calculated four SWR metrics: ripple duration, peak ripple frequency, peak ripple power, and sharp-wave amplitude. In 3 of 4 animals, PSILOCYBIN extended for hours while SALINE1 and SALINE2 recordings were only 1 hour. We therefore limited our analysis of the PSILOCYBIN sessions to the first hour following injection to match the length of the control SALINE1/SALINE2 sessions. We observed an acute increase in SWR duration during PSILOCYBIN compared to SALINE1 and SALINE2 (**Figure 2B, C**) which was consistent across all animals. We also observed a consistent decrease in peak ripple frequency across all animals during PSILOCYBIN compared to SALINE1/SALINE2 (**Figure 2E**). We observed an acute increase in sharp wave amplitude (**Figure 2F**) from SALINE1 to PSILOCYBIN across 3 of 4 animals. Finally, we observed a similar, acute increase in peak ripple power for 3 of 4 animals from SALINE1 to PSILOCYBIN (**Figure 2D**). These results held when we considered wake periods only (**Supplemental Figure 4**). Finally, we examined SWR features across the entire Psilocybin session in 30 minute blocks and found that while most features remained steady across time, ripple duration decreased for all four animals across the first hour (**Supplemental Figure 6C**). Nonetheless, we observed similar results when we examined the entirety of the PSILOCYBIN session, not just the first hour, though ripple duration was no longer significantly increased during the PSILOCYBIN session in one animal (**Supplemental Figure 5**). These results revealed that psilocybin acutely modulated SWR features, consistently producing longer, lower frequency ripples and increasing sharp wave amplitude in the majority of animals.

**Figure 2:**
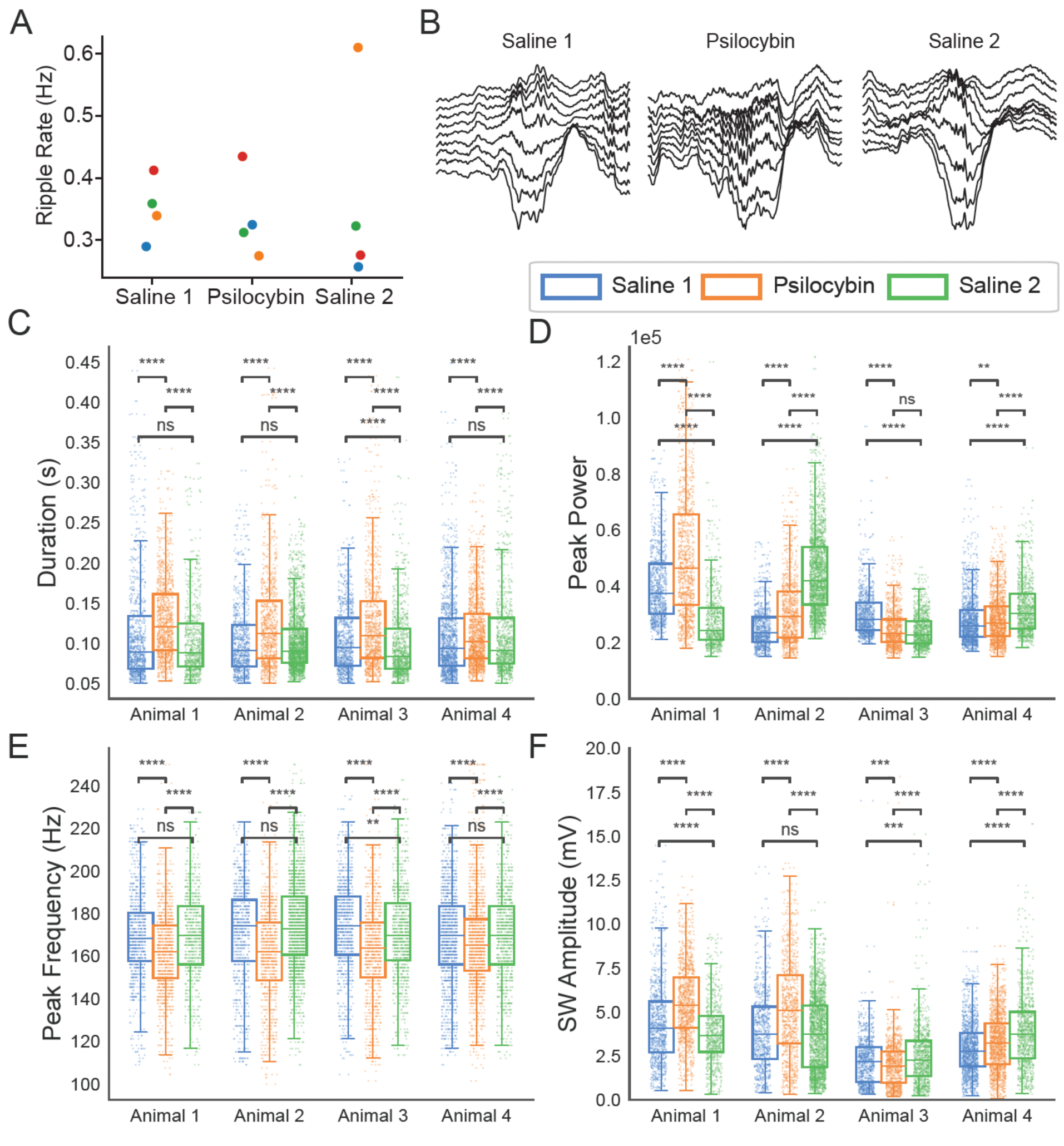
SWR features. **A)** Ripple rate for all animals across sessions. **B)** Example ripples of the median duration observed in each session. **C)** SWR duration for each rat shown across all three sessions shows a consistent increase in SWR duration under psilocybin. Only the first hour of data post-injection is shown. **D)** Same as C but for ripple power, which does not show consistent changes under psilocybin across animals. **E)** Same as C but for peak ripple frequency which shows a consistent decrease under psilocybin across all animals. **F)** Same as C but for sharp wave (SW) amplitude which shows a significant increase from Saline1 to Psilocybin sessions in 3 of 4 animals. Plots C) and F) are zoomed in for clarity. ****p<1e-4, ***p<0.001, **p<0.01, *p<0.05, two-sided Mann-Whitney U-test.

**Supplemental Figure 1.**
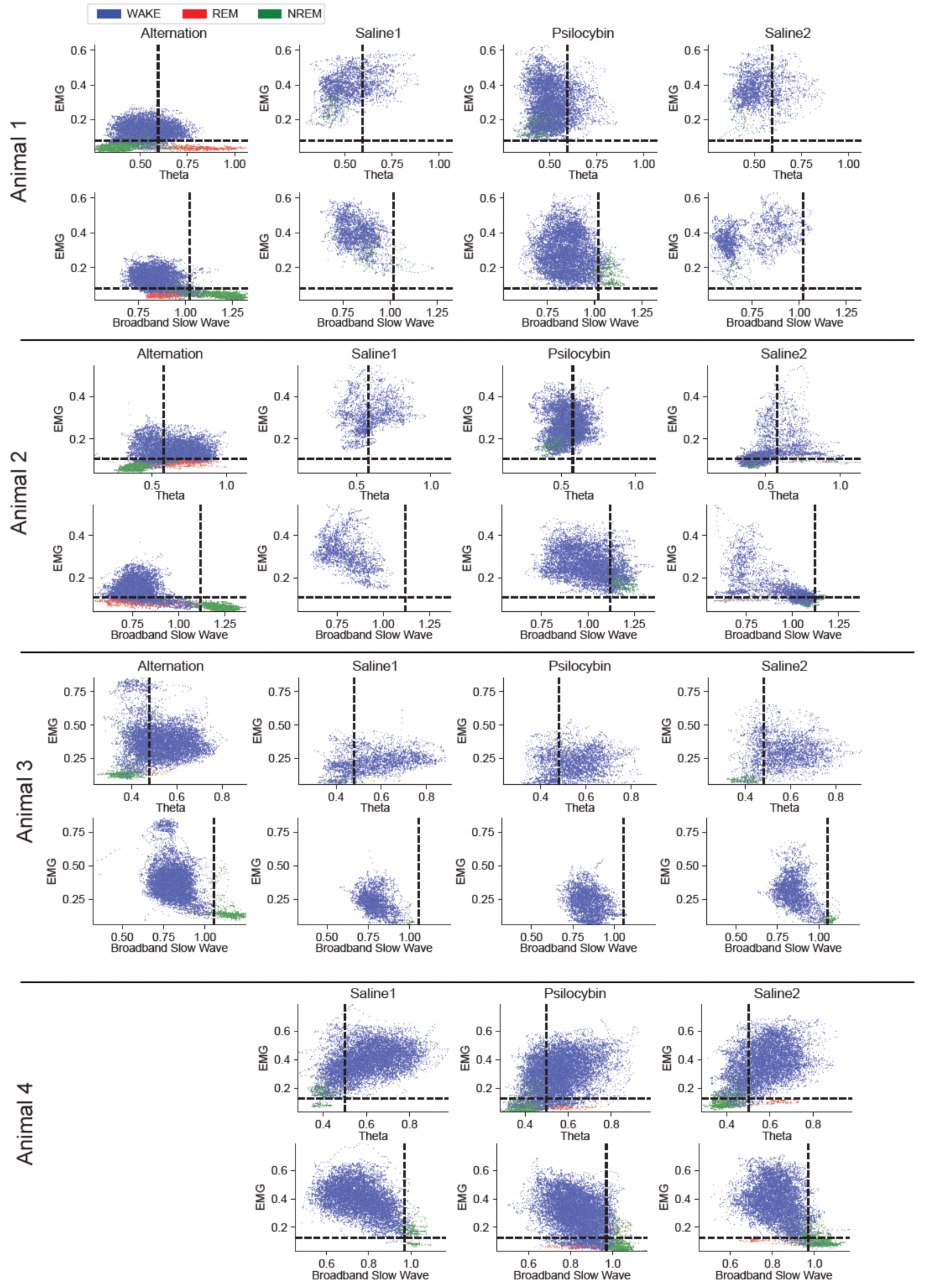
Sleep state features. Scatterplots of sleep scoring metrics in 1s bins for each animal across all sessions. Animals 1-3 exhibited little to no sleep in the SALINE1 and PSILOCYBIN sessions, so the alternation session used to establish thresholds for separating states is also shown on the left. For Animal 4, the Saline2 session exhibited sufficient sleep to establish thresholds and therefore no alternation session was necessary.

**Supplemental Figure 2.**
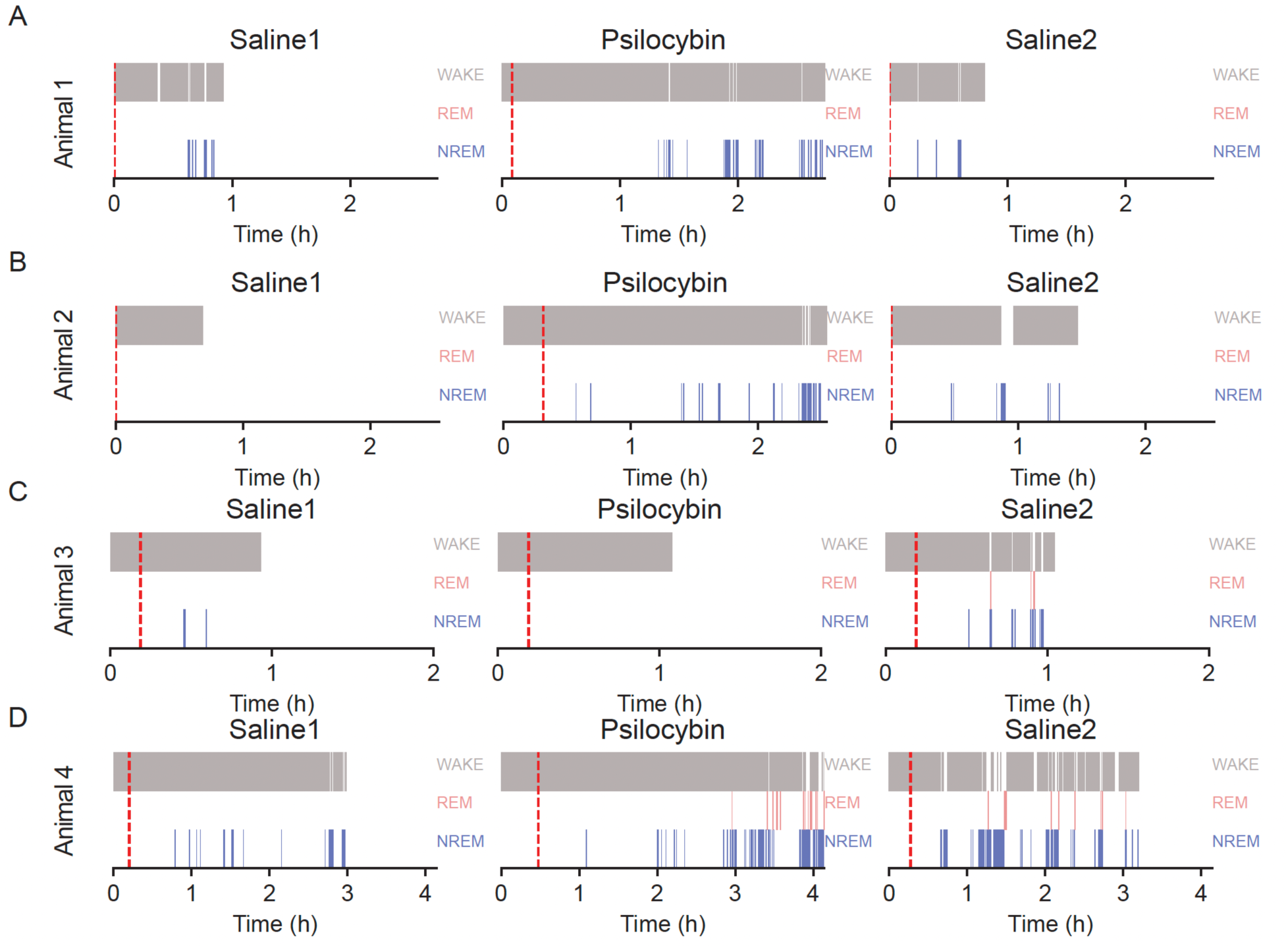
Hypnograms. **A)** Hypnogram showing periods of WAKE, REM, and NREM across all recordings for Animal 1. **B-D)** Same as A) but for Animals 2-4.

**Supplemental Figure 3.**
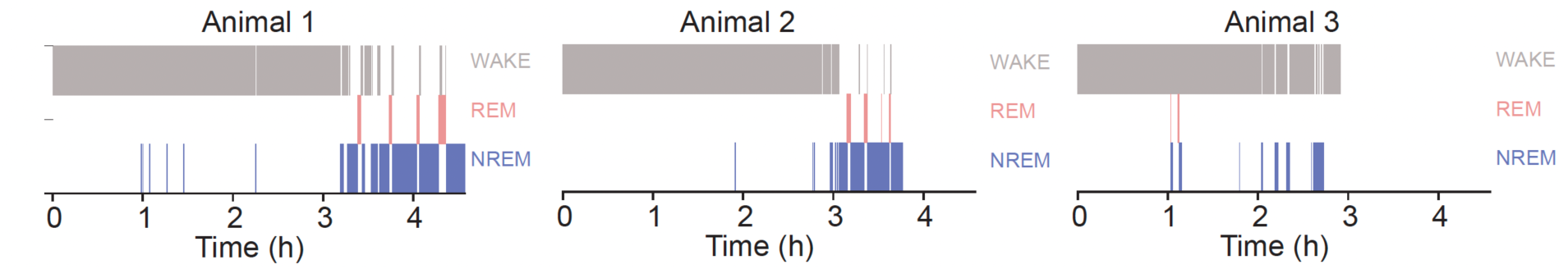
Hypnograms from Alternation session. **A)** Hypnograms showing periods of WAKE, REM, and NREM across all recordings for Animals 1-3 after the third consecutive day of learning a delayed spatial alternation task.

**Supplemental Figure 4.**
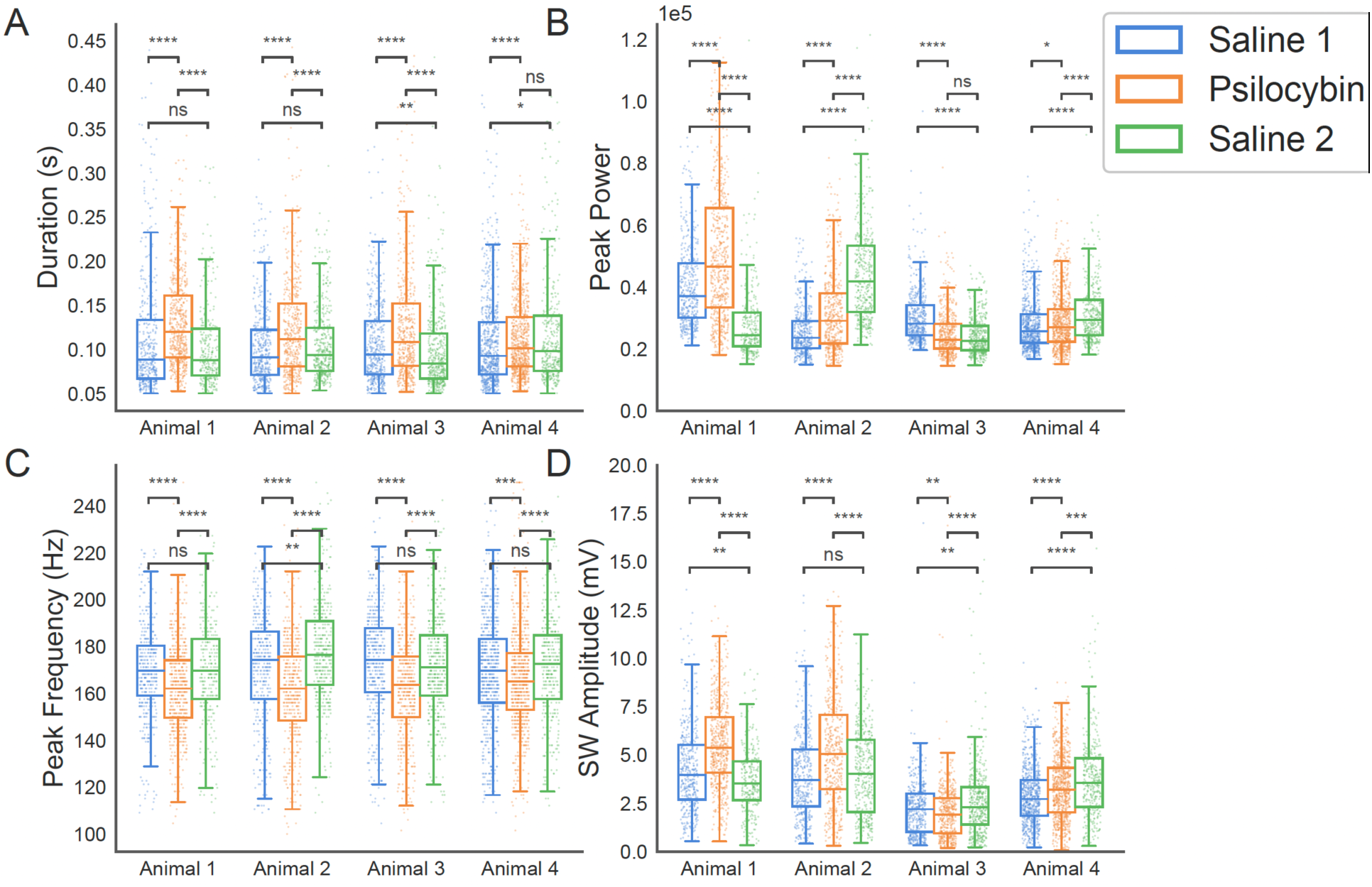
SWR features during wake states only. **A)** SWR duration for each rat shown across all three sessions shows a consistent increase in SWR duration under psilocybin during wake states only. Only the first hour of data post-injection is shown. **B)** Same as A but for ripple power, which does not show consistent changes under psilocybin across animals. **C)** Same as A but for peak ripple frequency which shows a consistent decrease under psilocybin across all animals. **D)** Same as A but for sharp wave (SW) amplitude which shows a significant increase from Saline1 to Psilocybin sessions in 3 of 4 animals. Plots B) and D) are zoomed in for clarity. ****p<1e-4, ***p<0.001, **p<0.01, *p<0.05, two-sided Mann-Whitney U-test.

**Supplemental Figure 5.**
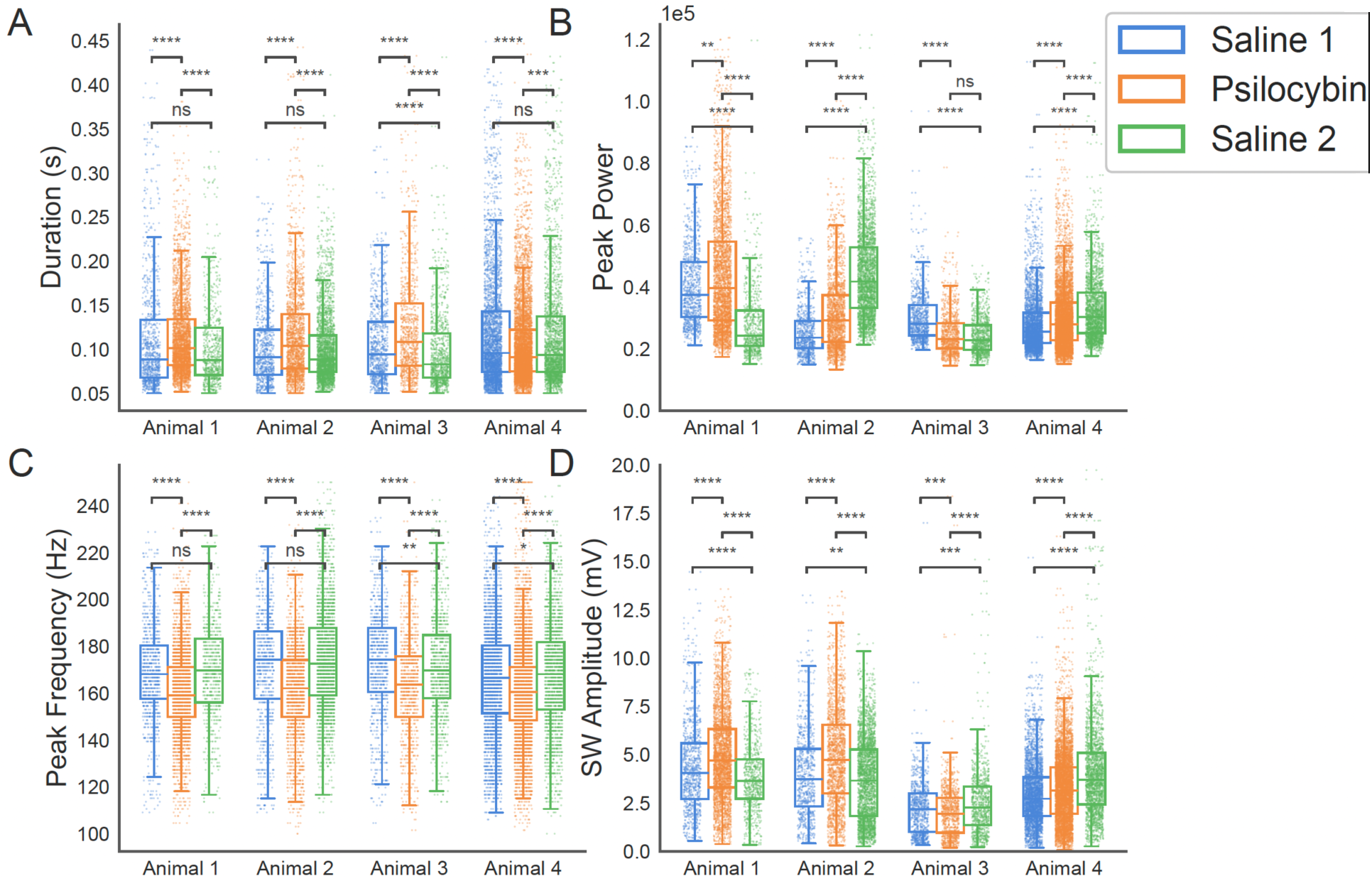
SWR features across entire session. **A)** Increased SWR duration during psilocybin vs Saline1 and Saline2 for animals 1-3. **B)** Ripple peak power showed animal-dependent changes rather than a consistent psilocybin effect. **C)** Decreased peak ripple frequency during Psilocybin vs Saline 1 and Saline 2 across animals. **D)** Increased sharp wave amplitude from Saline1 to Psilocybin session in Animals 1, 2, and 4. All available post-injection data were included, without restricting the analysis to the first hour. Panels B and D are zoomed for clarity. ****p < 1e-4, ***p < 0.001, **p < 0.01, *p < 0.05; two-sided Mann-Whitney U-test.

**Supplemental Figure 6.**
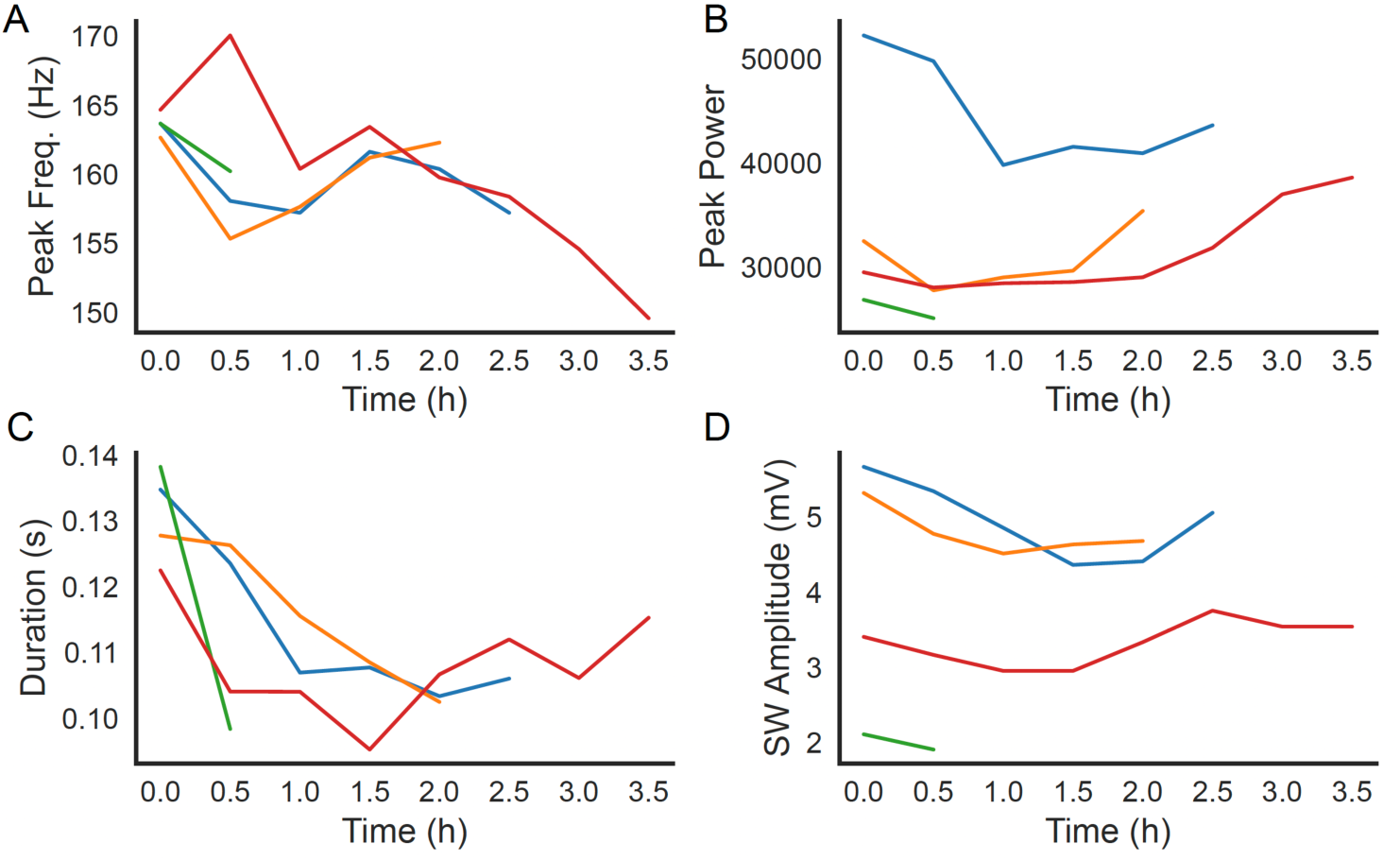
SWR features across time. **A)** Peak frequency for all four animals plotted over time in 30 minute bins across the entire post-injection Psilocybin session. **B)** Same as A but for peak ripple power. **C)** Same as A but for ripple duration showing a consistent drop in duration over the first hour post-injection. **D)** Same as A but for sharp wave amplitude.

## Discussion

In this study, we investigated how psilocybin impacts hippocampal sharp wave ripples. We found that that psilocybin produces a unique brain state in the hippocampus characterized by a high prevalence of HVS which are normally associated with transitions from wake to NREM sleep^48^. Despite observing reduced locomotion under psilocybin, however, we observed few such transitions from HVS to NREM. Psilocybin therefore produces a wake-like state which occurs prior to sleep but without transitioning into NREM. This finding is consistent with recent studies in rodents showing that lysergic acid diethylamide, a different psychedelic which also acts as a 5-HT2A receptor agonist, also increased HVS incidence^50^, and that psilocybin produced an acute decrease in NREM sleep^49^. These findings are also roughly consistent with the precursory body of knowledge on daytime psilocybin administration in humans where both delayed REM onset and REM duration were observed^51^. We note that animals also exhibited little sleep during the SALINE1 session. Though all animals were habituated to the general procedure of restraint for days prior to recording, SALINE1 was the first day they received the full injection. We therefore attribute low levels of sleep during SALINE1 to the novelty of the recording procedure and heightened stress following the i.p. saline injection since animals exhibited high levels of NREM and low NREM latency during SALINE2.

The sleep patterns observed during Saline2 could represent a return to normal sleep architecture following the novelty of injections and the psychedelic experience. However, sleep pressure can be exacerbated following drug induced wake periods, resulting in compensatory increased sleep duration in rodents^52^. Indeed, systemic administration of the serotonin agonist Quipazine has been shown to induce an initial period of sleep suppression^53^, even in previously sleep deprived animals^54^, followed by a rebound in NREM sleep. Thus, the observed rebound in NREM sleep during Saline2 could also indicate a recovery from an extended, psychedelic drug induced wake period.

We found that psilocybin robustly decreased ripple frequency and elongated ripple duration across all animals during the PSILOCYIN session. Ripple frequency is modulated by CA1 excitatory cell interactions with local interneurons^40^. Ripple duration is related to learning as long duration ripples are upregulated following a novel experience or after performing a memory task. Moreover, optogenetic ripple prolongation enhances memory and neuron firing patterns during longer ripples replay more diverse experiences, providing a causal link to memory^41^. We have previously shown that lower frequency, longer duration ripples also occur during sleep but not during acute sleep loss following a novel experience^55^, suggesting that psilocybin might mimic the beneficial effects of sleep. Furthermore, psilocybin could strengthen memory by linking discrete parts of a maze into a holistic spatial memory representation during replays which occur with long duration ripples. This would potentially enhance memory for the experience occurring immediately prior to psilocybin administration or for the psilocybin experience itself and would likely connect the pre- and post-psilocybin experiences together via recruitment into the same cell ensemble. Retrieval of one experience would therefore likely elicit memories of the other experience and explain why humans studies report a link between the positivity of the psychedelic experience, the context in which psilocybin is taken, and the subsequent therapeutic effects^56,57^. Future studies could investigate the effects of psilocybin on replay content and length. A previous study showed that the psychedelic lysergic acid diethtylamide produced a similar decrease in ripple frequency^50^. Unlike our results, this study also showed a large decrease in ripple rate and a small (∼5ms) decrease in ripple duration. Thus, perhaps due to differences in chemical structure^58^, different psychedelics produces distinct effects to SWRs.

We also observed an increase in sharp wave amplitude under psilocybin, though with more inter-animal variability. Sharp waves result when synchronous discharges of CA3 pyramidal cells depolarize downstream CA1 pyramidal cell dendrites located in *stratum radiatum^26,38^*. This increase in sharp wave amplitude suggests that psilocybin induces acute plasticity the CA3◊CA1 circuit. Future studies could examine whether the observed increase to sharp wave amplitude results from enhanced synaptic transmission in the CA3◊CA1 circuit or from more synchronous, longer lasting spiking input in CA3.

Longer ripples have been shown to produce strong reactivation of ensembles in prefrontal cortex^59^. Our results therefore indicate that psilocybin could increase functional coupling between the hippocampus and cortex, especially since learning-related coordination between hippocampal and prefrontal ensembles is stronger during wake than sleep^60^. By inducing longer ripples, psilocybin produces a state wherein the hippocampus is primed to interact with and induce plasticity in the cortex to produce stronger memories without inducing sleep. Enhanced memory for the psychedelic and immediately preceding experiences could therefore help decrease depressive symptoms by breaking out of excessive rumination on past negative experiences^61^. This effect is consistent with the “relaxed beliefs under psychedelics” model which holds that psychedelics’ therapeutic effects stem from their ability to disrupt established beliefs or “priors” which facilitates establishing new beliefs and priors about the world^62^. Thus, longer ripples could provide a neural mechanism underlying psilocybin’s therapeutic effects.

Many human studies using fMRI have found that connectivity within the default mode network decreases while brain-wide functional connectivity increases following psilocybin administration^63^. Drawing parallels with our results is difficult, however, since hippocampal engagement with the default mode network varies based on position along the anterior-posterior hippocampal axis (ventral-dorsal in rodents) as well as by subregion^64^ and many fMRI studies consider the hippocampus as one region. Our study recorded activity from the intermediate hippocampus regions CA1 which has widespread projections throughout the cortex including the prefrontal cortex^19^. However, this connectivity varies along the long axis of the hippocampus, with dorsal CA1 (posterior in humans) projecting to retrosplenial cortex and ventral CA1 (anterior in humans) projecting strongly to the amygdala, auditory and olfactory cortex, and thalamic / hypothalamic nuclei^18,19^. Additionally, activation of hippocampal projections from CA1 has been shown to inhibit the hypothalamic-pituitary axis to reduce stress-related glucocorticoid levels^25^. Thus, some of psilocybin’s therapeutic effects could derive from inhibition of the hypothalamic-pituitary axis by the hippocampus. Future studies could investigate whether psilocybin exerts a uniform or differential effect on ripples along the hippocampal long axis.

Overall, this study’s results indicate that psilocybin produces a unique brain state and induce longer, lower frequency ripples in the hippocampus. Future studies could investigate how hippocampal coupling with cortical regions changes following psilocybin administration, and how psilocybin modulates the content of hippocampal replays of recent and remote experiences which are linked to memory consolidation.

## Methods

### Animals

Male (n=2) and female (n=2) Long-Evans rats 5-12 months old were used in this study. Rats were socially housed in a vivarium on a 12 hour light-dark cycle with 1-2 other rats prior to surgery and were housed singly thereafter. Mice were given free access to food and water throughout the study. All procedures were performed in compliance with the guidelines of the University of Michigan Animal Care and Use Committee.

### Viral Constructs

For rat experiments, we used an pGP.AAV9.*Syn*.GcaMP7f.WPRE.SV40 virus from the University of Pennsylvania Vector Core/Addgene with an initial titer of 2.6x10^13^ GC/mL and diluted it into sterilized phosphate buffered saline (PBS) to a final titer of 2.6x10^12^ GC/mL for injection. Data collected using this virus was not examined in this study.

### Surgical procedures

We performed two stereotactic surgeries on naïve rats according to previously published procedures^65-67^. Both surgeries were performed under 1-2% isoflurane mixed with oxygen. Rats were given 2 mg/kg Meloxicam for analgesia orally prior to surgery and for two days following surgery. Animals were monitored daily for a minimum of seven days during recovery. Following induction of isoflurane anesthesia, a cocktail of a lidocaine (5mg/kg) and bupivacaine (2.5 mg/kg) cocktail was injected under the scalp to provide local anesthesia at the incision site. In the first surgery, 1000nL of GcaMP7f virus was infused in the prelimbic cortex at the center of a 1mm craniotomy (AP + 2.9, ML + 1.6, from Bregma, DV -3.0 at an 18 degree angle from top of brain). Following infusion, ∼1.5 mm of overlying cortex was removed, and a 23ga needle was lowered to ∼500µm above the target site. Then, a 0.6 x 7 mm GRIN lens was lowered to 3.0mm below the top of the brain, the area between the skull and lens was sealed with Kwik-Sil, and the lens was affixed to the skull with Vivid-Flow light-curable composite (Pearson Dental) and Metabond (Parkell). The lens was then covered in Kwik-Sil for protection. During this surgery, ground and reference screws were also placed over the cerebellum and a 3d printed crown base was attached to the rat’s skull^68^ to which crown walls and top were connected and to further protect the lens and future microdrive/probe implant.

8-16 weeks later, the rat was again given pre-operative Meloxicam and anesthetized under isoflurane. The crown walls were removed, copper mesh was attached to the crown base, and a customized V4 UCLA Miniscope with a short power cable with a connector attached to two header pins was permanently affixed to the skull to visualize calcium dynamics (not examined in this study). A 1.0mm craniotomy was then performed at AP-4.8, ML+3.6 from bregma. After removing dura and stopping bleeding with cold, sterile saline, a NeuroNexus A1x32-5mm-50-177 probe, attached to a metal microdrive, was implanted at 2.3 mm below the brain surface and the metal drive base was attached to the skull with Unifast light cured dental epoxy (Henry-Schein). The craniotomy was sealed with Dow-Sil, the probe was protected with dental wax, and the ground and reference wires were connected to the probe electronic interface board (EIB). A custom protective crown was then constructed from copper mesh, the EIB and miniscope power connector were connected to the crown walls, and the rat was removed from isoflurane and allowed to recover. The rat was monitored daily for 7 days prior to recording, during which the probe was lowered ∼1mm until ripples and spiking activity were visualized on the middle electrodes with sharp waves on at least 5 electrodes below indicating localization of the probe in the middle of the CA1 cell layer.

### Recording procedure

Recordings occurred across three consecutive days in three of four animals. In one animal there was a two-day gap between the Psilocybin and Saline2 session. Prior to psilocybin injections, electrophysiological, imaging, and behavioral video data were recorded for 5-15 minutes (PRE) in a rest box identical to the animal’s home cage but with an open top. PRE recordings were also included for saline sessions in two animals. For one animal, bedding from another animal’s cage (former cage mate) was placed in the bottom of the rest box for all sessions. In the other animals, a white towel was placed in the rest box. Following the PRE recording, the animal was disconnected and given an I.P. injection of either psilocybin (10 mg / kg) or an equivalent amount of saline. Recording cables were then re-connected and the animal was placed back in the rest box for a post-injection recording (POST) which lasted 1 hour following saline injections and 3-4 hours following psilocybin injections. In one animal the psilocybin recording was stopped at 1 hour post injection due to increased motion inducing excessive disconnects of the recording cable at that time. In another animal, POST recordings following saline injection were extended to 4 hours to match the psilocybin sessions. Since this animal exhibited extended periods of sleep during the Saline2 session, that session was used as a baseline for establishing thresholds for sleep scoring. The other three animals also performed a delayed spatial alternation task several weeks prior across five consecutive days. In this task, rats were first water deprived and then had to traverse the center stem of the T-maze where they freely chose to turn left or right and turn down the return arms back to the center stem start where they were blocked in for 10 seconds before starting another trial. Water was automatically delivered for either choice on the first trial following which water was only delivered if the opposite arm was visited. Following day 3, when each animal exhibited a jump in performance exhibited by > 75% correct trials and at least 10 correct trials in a row, the animals were placed in the rest box and neural activity was recorded for at least 4 hours. Extended post-alternation recordings were not performed on other days and only the post-performance sleep / rest data is analyzed here.

### Data acquisition and processing

Electrophysiological data were obtained using an Intan 1024ch Recording Controller using OpenEphys software at a sampling rate of 30,000Hz. The UCLA Miniscope utilizes a high amplitude alternating voltage (∼ 4800Hz) to modulate the electrowetting lens (EWL) shape for adjusting the imaging focal plan which infiltrates the signal recorded by the nearby silicon probe electrodes. Therefore, prior to performing any analyses, this noise was removed using a series of notch filters at the EWL holding frequency and its harmonics. De-noised data was then downsampled to 1250 Hz for subsequent local field potential analyses.

### Sleep state detection

Sleep states were scored in 1 second bins in line with previous studies^69,70^ using open-source code (the SleepScoreMaster function) adapted from https://github.com/buzsakilab/buzcode. Briefly, we first calculated three variables for sleep scoring: 1) pseudo-EMG (pEMG) calculated from the summed pairwise correlations between all electrodes after applying a 300-600Hz bandpass filter^71^, 2) theta power, defined as the maximum power in the 5-10Hz range above the aperiodic component of the power spectrum, and 3) broadband slow-wave power, defined as the slope of the periodic component^72,73^ of the power spectrum in the 4-90 Hz range^74^. For each animal, thresholds were determined by aggregating values for all metrics across an extended non-psilocybin session (Supplemental Figure 1) at least 3 hours long (a post-alternation session for 3 animals, and Saline2 session for the 4^th^ animal) and setting the threshold as the observed bimodality in the data. Setting thresholds from different sessions for each animal was necessary because we observed no sleep (and therefore no bimodalities in pEMG, theta power, and broadband slow-wave power) in the PSILOCYBIN sessions. Remaining periods with high theta and low pEMG were designated as REM. Wake time were designated as all remaining times. Finally, all sleep states were visually inspected and manually curated to adjust state transition times.

#### High Voltage Spindle detection

High voltage spindles (HVS)^47^ were detected by filtering signal from an electrode near the hippocampal pyramidal layer in the HVS band (4-9Hz) and its first harmonic (10-20Hz), identifying candidate periods of high power in both frequency bands (thresholds: HVS = 4 std, harmonic = 2 std), extending these candidate periods to when the power in each band fell below 1 std, and then defining their intersection as an HVS epoch.

#### SWR detection and analysis

First, we identified the channel closest to the pyramidal cell layer through visual inspection as the electrode where the deflection during each sharp wave changed from positive to negative. Second, periods with large voltage deflections were detected and excluded from further analysis. Third, ripple-band activity was isolated by applying a 150-250 Hz bandpass filter to local field potential, smoothed with a 125ms gaussian kernel, squared to produce a power envelope, and z-scored. Periods with z-scored ripple-band power > 4 for at least 50 ms were identified as candidate ripple events^39^. The start and end of each candidate ripple were then extended to the time when the z-scored power crossed below 0.5.

A wavelet analysis was then performed to identify the peak power and peak frequency of each ripple^75,76^. We next extracted signal from the pyramidal layer channel and the four neighboring electrodes above and below it (50 µm spacing), bandpass filtered these signals in the 2-30Hz range and calculated sharp wave amplitude as the maximum (peak-to-peak) difference between the bandpass filtered voltage observed across all channels during each ripple. Code used to identify SWRs and quantify each SWR feature is available at www.github.com/diba-lab/NeuroPy

#### Statistics

Unless otherwise noted, condition-wise differences between Saline1, Psilocybin, and Saline2 sessions were tested using a two-sided Mann-Whitney U-test. Figures used boxplots with overlaid points and conventional significance annotations.

## Acknowledgments

We would like to thank Hazel Jackson for detailed feedback on this manuscript. Ilknur Kayikcioglu Bozkir was supported by the Scientific and Technological Research Council of Türkiye (TÜBİTAK) under the 2219 International Postdoctoral Research Fellowship Program.

## Author Contributions

Conceptualization: N.R.K and K.D. Methodology: N.R.K. and K.D. Software: N.R.K., I.B., R.L. Validation: N.K., I.B., R.L. Formal Analysis: N.R.K., I.B., R.L. Investigation: N.R.K., T. L. Resources: K.D., D.P. Data Curation: N.R.K., I.B., R.L. Writing – original draft preparation: N.R.K., I.B., R.L. Writing – review and editing: N.R.K., I.B., R.L., D.P. Visualization: N.R.K., I.B., R.L. Supervision: N.R.K., K.D. Project administration: N.R.K. Funding Acquisition: N.R.K., K.D., D.P.

